# A bispecific immunotweezer prevents soluble PrP oligomers and abolishes prion toxicity

**DOI:** 10.1101/386763

**Authors:** Marco Bardelli, Karl Frontzek, Luca Simonelli, Simone Hornemann, Mattia Pedotti, Federica Mazzola, Manfredi Carta, Valeria Eckhardt, Rocco D’Antuono, Tommaso Virgilio, Santiago F. González, Adriano Aguzzi, Luca Varani

## Abstract

Antibodies to the prion protein, PrP, represent a promising therapeutic approach against prion diseases but the neurotoxicity of certain anti-PrP antibodies has caused concern. Here we describe scPOM-bi, a bispecific antibody designed to function as a molecular prion tweezer. scPOM-bi combines the complementarity-determining regions of the neurotoxic antibody POM1 and the neuroprotective POM2, which bind the globular domain (GD) and flexible tail (FT) respectively. We found that scPOM-bi confers protection to prion-infected organotypic cerebellar slices even when prion pathology is already conspicuous. Moreover, scPOM-bi prevents the formation of soluble oligomers that correlate with neurotoxic PrP species. Simultaneous targeting of both GD and FT was more effective than concomitant treatment with the individual molecules or targeting the tail alone, possibly by preventing the GD from entering a toxic-prone state. We conclude that simultaneous binding of the GD and flexible tail of PrP results in strong protection from prion neurotoxicity and may represent a promising strategy for anti-prion immunotherapy.

**Author summary:** Antibody immunotherapy is considered a viable strategy against prion disease. We previously showed that antibodies against the so-called globular domain of Prion Protein (PrP) can cause PrP dependent neurotoxicity; this does not happen for antibodies against the flexible tail of PrP, which therefore ought to be preferred for therapy.

Here we show that simultaneous targeting of both globular domain and flexible tail by a bispecific, combination of a toxic and a non-toxic antibody, results in stronger protection against prion toxicity, even if the bispecific is administered when prion pathology is already conspicuous.

We hypothesize that neurotoxicity arises from binding to specific “toxicity triggering sites” in the globular domain. We designed our bispecific with two aims: i) occupying one such site and preventing prion or other factors from docking to it and ii) binding to the flexible tail to engage the region of PrP necessary for neurotoxicity.

We also show that neurotoxic antibodies cause the formation of soluble PrP oligomers that cause toxicity on PrP expressing cell lines; these are not formed in the presence of prion protective antibodies. We suggest that these soluble species might play a role in prion toxicity, similarly to what is generally agreed to happen in other neurodegenerative disorders.

## Introduction

Prions are the causative agent of sporadic, hereditary and iatrogenic forms of transmissible spongiform encephalopathies, which afflict humans and broad spectrum of mammals and are invariably fatal[1–3]. Whereas Bovine Spongiform Encephalopathy (the most prevalent prion disease in the 1990s, also known as “Mad Cow” disease) has been largely defeated, Chronic Wasting Disease, which affects deer, elk and moose, remains prominent in parts of the US, Canada, South Korea and has recently reached Norway[4]. These findings are raising renewed concerns about the contamination of the food chain. Transmission of infectious animal material to humans causes variant Creutzfeldt-Jakob disease (vCJD). A common Met/Val polymorphism at codon 129 of the *PRNP* gene is assumed to be important in susceptibility of humans to prion infections, with homozygous individuals (Met/Met and Val/Val) being overrepresented in collectives of CJD patients[5]. The recent discovery of a case of vCJD in a 36-year-old man producing both M129 and V129 variants of PrP, which is much more frequent in the population and is thought to conduce to a disease developing more slowly, has led to suggestion that we might be facing a new wave of vCJD cases [6].

Human prion diseases continue to be intractable and are poorly understood at the molecular level. It is firmly established that conversion of cellular prion protein (PrP^C^) into a toxic, self-replicating form (scrapie, PrP^Sc^) leads to the formation of aggregates [2,7]. How such aggregates induce toxicity, however, is largely unknown.

Antibodies against PrP have been proposed as a valid therapeutic strategy against prion diseases, much like antibodies targeting the amyloid-β protein are showing promise in clinical trials against Alzheimer’s disease (AD) [8,9]. Furthermore, anti-PrP antibodies were shown to be protective in preclinical models of AD [10]. On the other hand, recent literature reports [11] have highlighted potential safety issues, since an epitope-dependent subset of anti-PrP antibodies have been found to cause prion mediated neurodegeneration.

PrP has a structured globular domain (GD), whose general architecture is highly conserved amongst mammals, and an unstructured N-terminal region, often referred to as flexible tail (FT). Several antibodies against the globular domain were shown to elicit neuronal toxicity [11,12]. The exact epitope on the GD, rather than the binding affinity or other properties, appears to be the main determinant of toxicity. One such neurotoxic antibody with intriguing properties, POM1, was extensively characterized [11,13]. POM1 binds with low nanomolar affinity to a discontinuous epitope comprising the α1-α3 region of the GD. PrP expressing mice and cerebellar organotypic cultured slices (COCS) exposed to POM1 show rapid neurotoxicity [11]. Intriguingly, toxicity was prevented by a deletion of the octapeptide repeats in the FT, even if this deletion did not affect the POM1 epitope. Antibodies against the FT, such as POM2, were also capable of preventing POM1-mediated toxicity, but only if administered before POM1.

POM1 and *bona fide* prions exert similar toxic effects such as neuronal cell loss, astrogliosis, microgliosis and spongiform change. PrP^Sc^, the proteinase K-resistant form of prion protein, is used as a surrogate marker for prions and believed to be the infectious agent[14]. In both POM1 and prions, metabotropic glutamate receptors play an important role in downstream toxicity and compounds that rescue POM1-induced toxicity also alleviate PrP^Sc^-induced toxicity [15,16].

Overall, the observations indicate that antibody binding to specific sites in the globular domain triggers a toxic process, mediated by the FT, which converges with that initiated by infectious prions. However, toxic prion antibodies do not generate prion infectivity [17]. The above leads us to propose that the binding of POM1 to a “toxicity triggering site” might emulate the docking of PrP^Sc^ (or other toxic factors) to the GD. According to this hypothesis, a molecule that occupies the POM1 binding site in the GD might prove beneficial by preventing interaction with PrP^Sc^ and, consequently, toxicity.

Here we designed a bispecific antibody formed by the POM1 variable region, which binds the GD, and the POM2 variable region, which binds to the FT and was shown to prevent POM1-mediated toxicity. We show here that the POM1-POM2 bispecific single chain antibody (called scPOM-bi), a combination of the toxic POM1 and non-toxic POM2 antibodies, is capable of preventing prion toxicity in COCS even when given 21 days post infection, when signs of prion pathology are already visible[18]. POM1 forms soluble PrP-containing oligomers that cause toxicity to PrP expressing cells. By contrast, delivery of scPOM-bi prevents formation of soluble oligomers. scPOM-bi shows increased protection in comparison to the isolated POM2 or to a mixture of POM1 and POM2, despite having similar binding affinity, suggesting that simultaneous targeting of globular domain and flexible tail might represent an optimized strategy for immunotherapy of prion diseases.

## Results

### Design of scPOM-bi, a bispecific POM1-POM2 antibody

We fused the toxic antibody POM1 with POM2, capable of preventing its toxicity if administered before POM1, by joining their variable regions in single chain format with a (GGGGS)_x3_ linker commonly utilized in non-natural antibodies [19,20]. We have had considerable success in using this format to produce bispecific antibodies against various targets. Although the order of the variable fragments and linker size might affect binding and efficacy, computational simulations showed that the chosen design, with the POM1 moiety preceding POM2 and the chains arranged as VL-VH-VH-VL, was compatible with binding to PrP. Indeed, the resulting bispecific nanobody, scPOM-bi, was produced in *E. coli* and characterized to be functional, monomeric and folded with a melting temperature of 75 °C (S1 Fig.). We did not, therefore, explore the production of alternative constructs. Computational docking and molecular dynamics simulations, based on the available experimental structures of POM1 in complex with PrP globular domain [21] and POM2 bound to a FT derived peptide [22], indicate that the size and orientation of scPOM-bi is compatible with bivalent engagement of PrP (Fig 1).

**Fig. 1:**
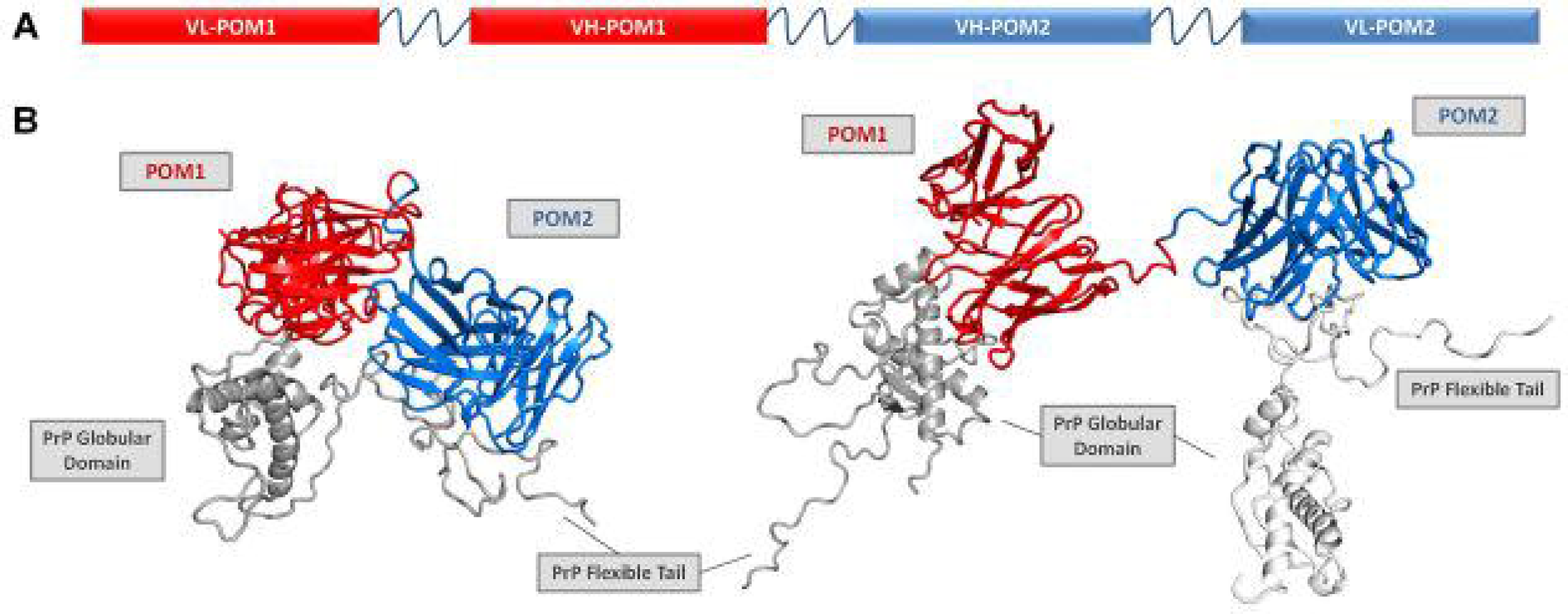
A bispecific immunotweezer formed by a combination of toxic (POM1) and non-toxic (POM2) antibodies. (A) The single chain variable domains of POM1 and POM2 are joined by a flexible linker to yield the bispecific scPOM-bi, schematic representation. B) Computational molecular dynamics model of scPOM-bi in complex with mPrP; intra-molecular (left and S1 movie) and inter-molecular (right and S2 movie) binding modes are shown. They are both structurally accessible to scPOM-bi.

### scPOM-bi protects from prion toxicity when administered late after prion infection

When infected with prions, COCS faithfully mimic prion pathology and are easily accessible to pharmacological manipulation [23]. In contrast to POM1, chronic treatment with scPOM-bi for 21 days did not produce observable toxicity in COCS despite comprising the toxic POM1 moiety (Fig 2A). Furthermore, simultaneous addition of the individuals POM1 and POM2 to COCS resulted in neurotoxicity, indicating that the bispecific has different properties than the simple sum of its parts. We added scPOM-bi to COCS from PrP^C^-overexpressing *Tga*20 mice infected with either RML6 prions (RML6 = passage 6 of the Rocky Mountain Laboratory strain, mouse-adapted scrapie prions) or non-infectious brain homogenate (NBH) as a control. 45 days post infection (dpi), immunohistochemical staining for NeuN, identifying neurons, showed widespread neuronal degeneration in the presence of RML but not NBH. By contrast, treatment with scPOM-bi prevented RML-induced neurotoxicity even when administered at 21 dpi (Fig 2B and S2 Fig). The anti-FT antibody POM2 did not afford similar protection levels at 21 dpi, despite being used at 5-fold higher molarity than scPOM-bi (Fig 2B) and having comparable binding affinity for PrP (Fig 3). PrP^Sc^ levels, detected by proteinase K-digestion of tissue inoculated with RML, remained constant in prion-infected *Tga20* COCS treated with scPOM-bi for 21days (Fig 2C), suggesting that neuroprotection is not primarily mediated by reduced amounts of PrP^Sc^ in RML infected COCS.

**Fig. 2:**
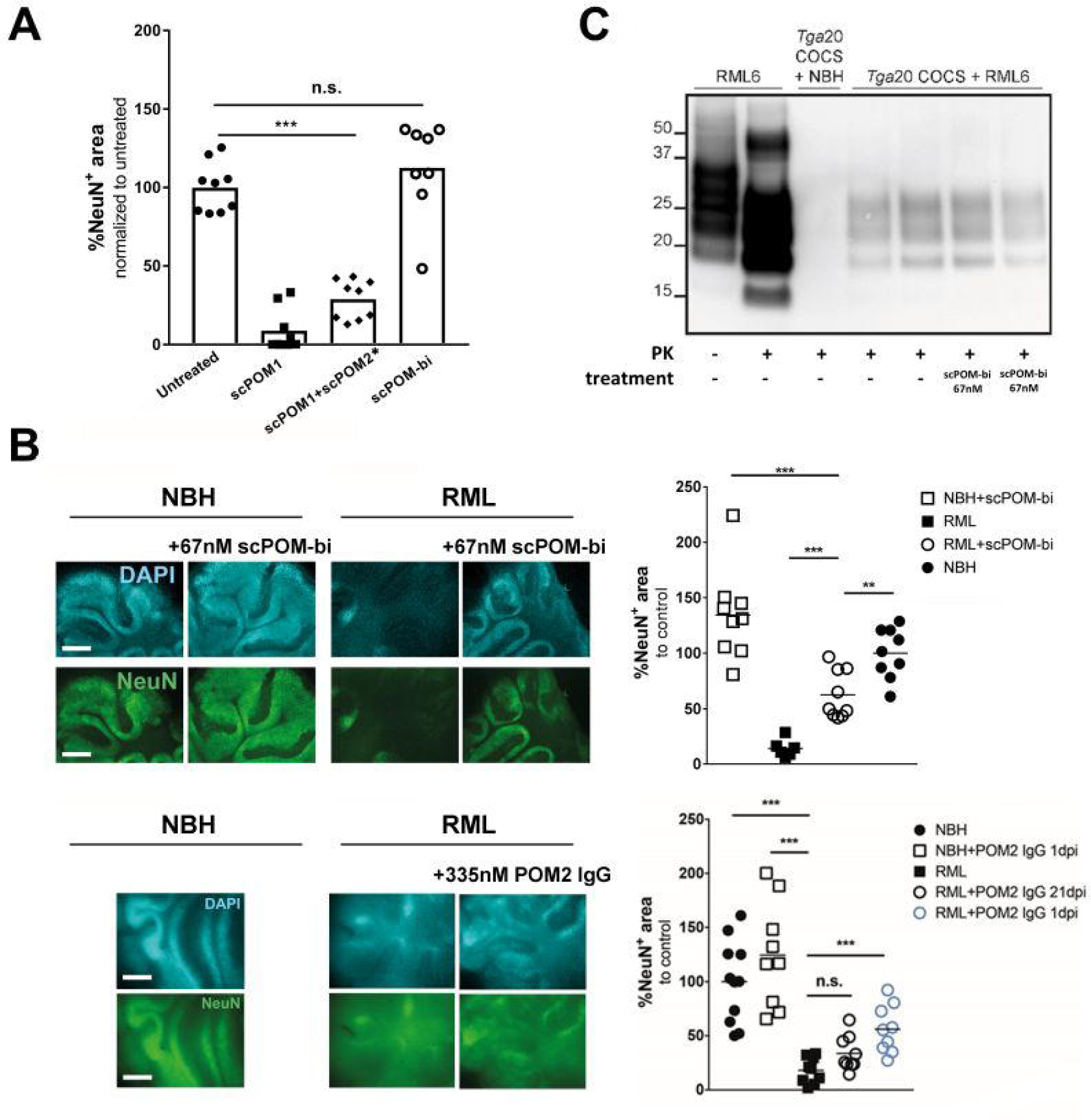
The bispecific scPOM-bi antibody protects against prion infection even when administered 21 days post infection (dpi). (A) Chronic treatment with scPOM-bi for 21 days did not produce observable toxicity in COCS, contrary to POM1. Furthermore, simultaneous addition of the individuals POM1 and POM2 to COCS resulted in neurotoxicity, indicating that the bispecific has different properties than the simple sum of its parts Area staining of neuronal nuclei by NeuN is shown on the y axis (lower values correlate with toxicity). Column 3 (*) is from a different experiment with a related negative control on which the data was normalized to. (B) scPOM-bi prevents RML induced neurotoxicity even when added 21 dpi (top). Despite similar binding affinity for PrP, POM2-IgG does not achieve similar protection at 21 dpi even at 5 fold higher concentration (bottom). COCS inoculated with non-infectious brain homogenate (NBH) are used as control; the images show NeuN and DAPI staining of COCS, scale bar = 500 µm. ** p<0.01, *** p<0.001, n.s. = not significant, one-way ANOVA with Dunnett’s post-hoc test. *Upper panel*: n=9 biological replicates (1 COCS = 1 biological replicate) for all treatment groups except for RML alone (n=8). *Lower panel*: n=9 biological replicates for all treatment groups. Images of all biological replicates depicted in S2Fig. (C) Western blot shows the presence of PK resistant material in COCS inoculated with RML. Addition of scPOM-bi 21 days after prion inoculation of Tga20 COCS did not show conceivable reduction of PrP^Sc^.

**Fig. 3:**
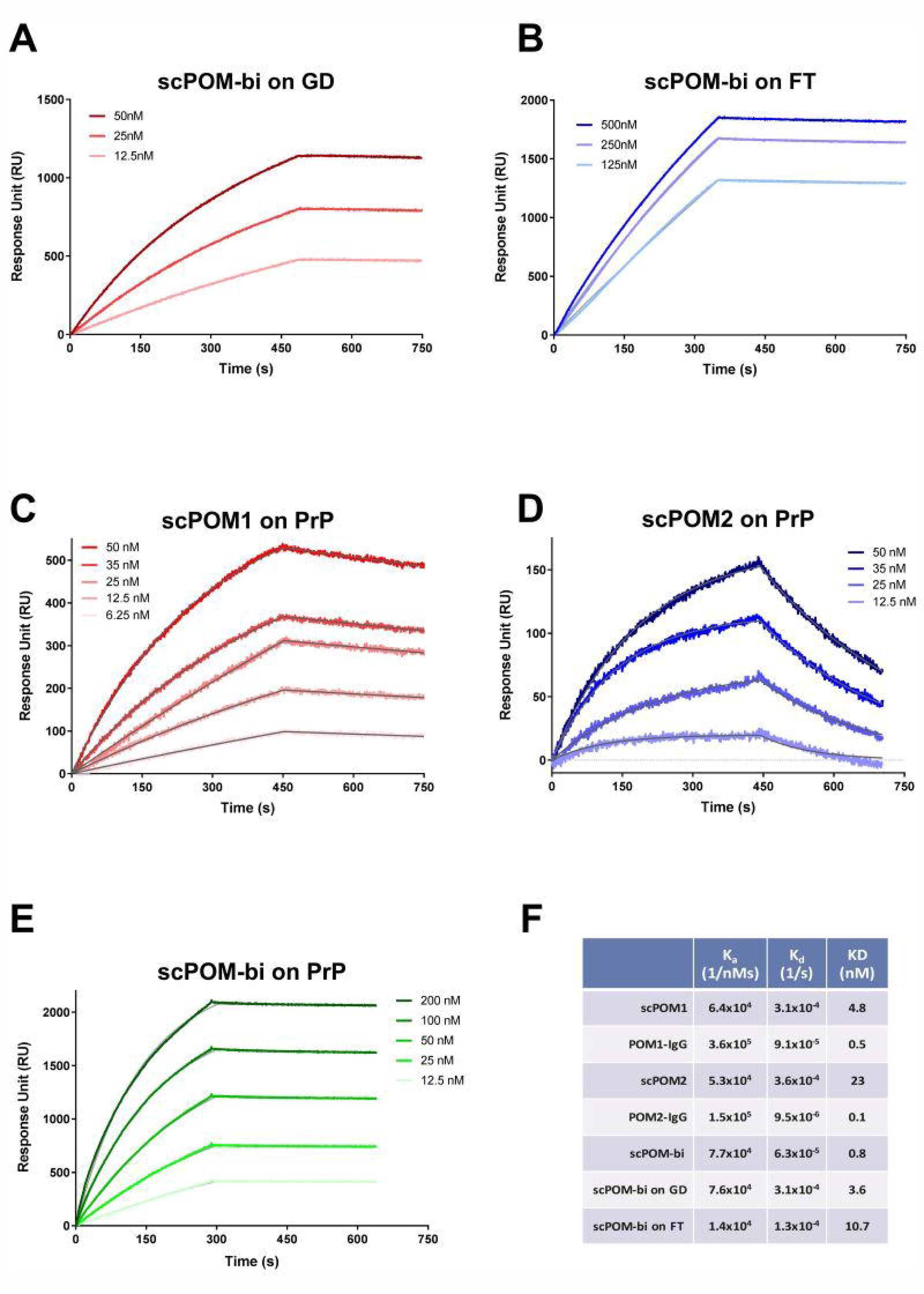
the bispecific antibody scPOM-bi binds simultaneously to GD and FT of PrP with high affinity. SPR sensorgrams for binding of scPOM-bi to truncated PrP constructs lacking FT (A) or GD (B) indicate that both antigen binding sites of scPOM-bi correctly engage their target. The bispecific antibody (E) had a stronger affinity than its individual components (C-D) due to avidity resulting in a slower dissociation. The fitting of the experimental data used to calculate the binding constants is in grey. Values for the above plus the full IgG versions of POM1 and POM2 are summarized in (F).

### scPOM-bi binds with high affinity to globular domain and flexible tail of PrP

Surface plasmon resonance (SPR) assays showed that scPOM-bi binds PrP with low nanomolar affinity, stronger than that of its individual single chain components due to avidity effects, resulting in slower dissociation, when the two antigen binding sites are engaged (Fig 3). Indeed, scPOM-bi was found to bind to both a PrP construct lacking the FT and to one lacking the GD with affinity similar to that of its individual components, scPOM1 and scPOM2 (Fig 3), confirming that both paratopes of scPOM-bi correctly recognize their cognate epitopes. scPOM-bi avidity can arise from two binding modes: simultaneous engagement of the GD and FT binding sites on a single PrP molecule (intramolecular) or binding of the POM1 site on one PrP molecule and the POM2 site on another (intermolecular). Both options are structurally allowed according to molecular dynamics simulations (Fig 1). SPR assays performed with different quantities of immobilized PrP indicate that intermolecular binding is available to scPOM-bi (see S1 Text and S3 Fig. for details).

### scPOM-bi inhibits the formation of partially PK resistant soluble PrP oligomers

POM1 binds the GD and was shown to have neurotoxic effects mediated by the FT [11]. Soluble oligomers may be the toxic species responsible for Alzheimer’s and other amyloidosis [24–28] and the smallest infectious unit of prions may also be oligomeric [29]. We thus used dynamic light scattering (DLS) to compare the aggregation properties of toxic and protective antibodies.

Single species of size compatible with the monomeric forms of uncomplexed recombinant mouse Prion Protein (mPrP) or antibodies were observed by DLS. Addition of the toxic antibody POM1 to mPrP *in vitro*, instead, caused the formation of soluble oligomers with a radius of approximately 200nm (Fig 4A). As a reference, the monomeric scPOM1:GD complex has an elliptical shape of ~7 × 5 nm according to the available x-ray structure[13]. The radius value is derived from interpretation of DLS data using a spherical model which may or may not be appropriate for these molecules. This, however, does not affect the following qualitative and comparative analysis of the results.

**Fig. 4:**
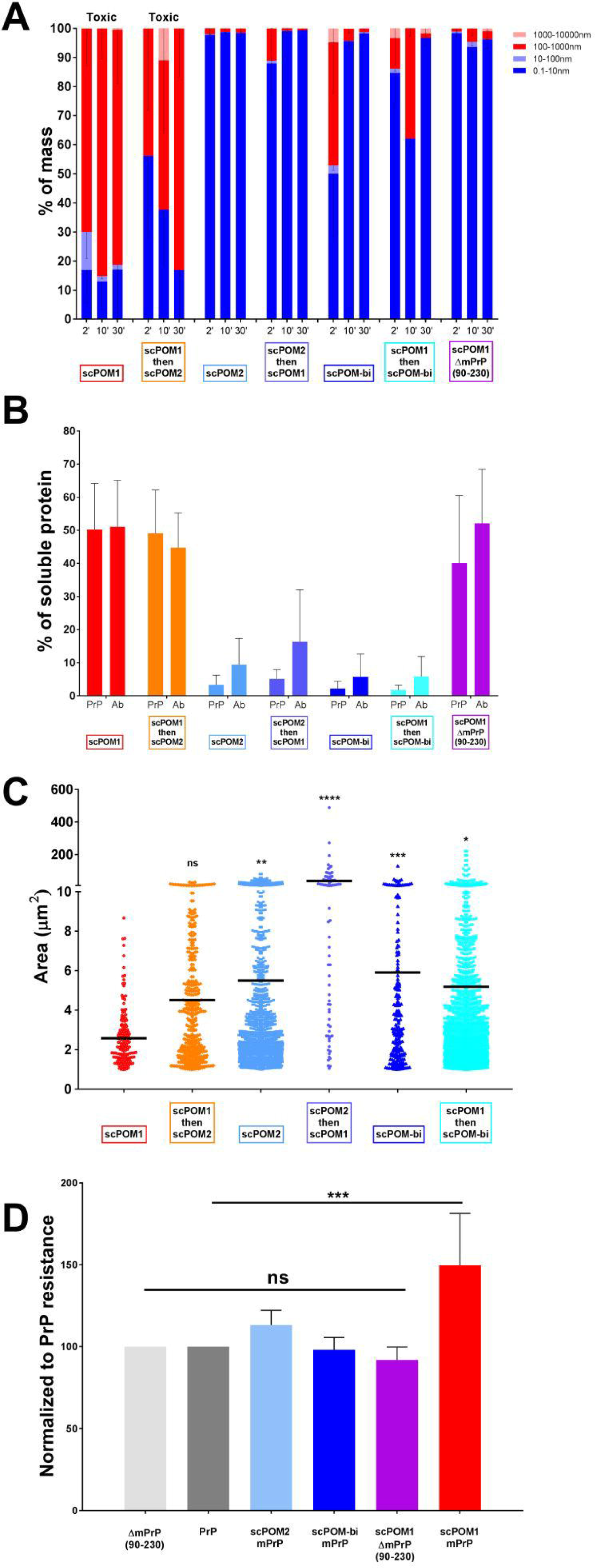
scPOM-bi prevents the formation of soluble, PK resistant oligomers. (A) DLS showed the presence of soluble oligomers (red shades in histograms, reported as percentage) upon addition of the POM1 toxic antibody to recombinant mPrP in vitro. Subsequent addition of POM2 did not remove the oligomers or inhibit toxicity. Smaller species comparable to monomeric forms (blue) were detected in solution when POM1 was in complex with ΔmPrP^C90-230^, lacking the flexible tail, and when POM2 was added to mPrP prior to POM1 addition. Similarly small species were found when the neuroprotective scPOM-bi was added to mPrP; the bispeficic was also capable of removing the soluble oligomers generated by POM1. DLS data is shown for 3 time points after complex formation. (n=5 for scPOM1:mPrP, scPOM2:mPrP and scPOM-bi:mPrP ; n=3 for scPOM1:mPrP then scPOM2, scPOM2:mPrP then scPOM1 and scPOM1:mPrP then scPOM-bi) (B) DLS can only detect soluble material. To investigate the presence of insoluble aggregates we formed the mPrP:Ab complexes *in vitro*, centrifuged them and analyzed the resulting supernatant with PAGE/Western blot. Soluble material was only detected in toxic combinations (POM1:mPrP or POM1:mPrP followed by POM2, red and orange). The percentage of mPrP and antibody in solution (normalized against isolated PrP or antibody) is shown; data from quantification of band intensity on SDS-PAGE (images in S5 Fig. – n=7 for all samples tested). (C) In order to characterize both soluble oligomers and insoluble aggregates we formed the mPrP:Ab complexes *in vitro* and deposited the resulting material on microscopy slides. Confocal microscopy indicates that toxic antibody combinations (e.g. POM1:mPrP or POM1:mPrP followed by POM2, red and orange) generate species with smaller average size than protective antibody combinations. The surface area of the detected species is reported on the y axis, the horizontal line represents the average. Differences can also be appreciated by visual inspection of the confocal microscopy images (S4 Fig – scPOM1:mPrP n=166, scPOM2:mPrP n=1136, scPOM-bi:mPrP n=204, scPOM1:mPrP then scPOM2 n=444, scPOM2:mPrP then scPOM1 n=74 and scPOM1:mPrP then scPOM-bi n=1767). D) The soluble oligomers generated by POM1 showed increased resistance to *in vitro* degradation by proteinase K at 2μg/ml (red). Such resistance was abolished when POM1 bound a mPrP construct lacking the FT (light red) or in non-toxic antibodies (shades of blue). Data from quantification of PK resistant bands on western blot, normalized against isolated PrP (images in S6 Fig. – scPOM1:mPrP n=5, scPOM1:ΔPrP n=3, scPOM2:mPrP n=4, scPOM-bi:mPrP n=4).

When POM1 was added *in vitro* to a truncated version of mPrP, ΔmPrP(residues 90-230), lacking the N-terminal FT, only monomeric species were found in solution. The disordered region of PrP, in other words, was required for the formation of POM1-induced soluble aggregates just like POM1-induced neurotoxicity is abrogated in the absence of the FT. Notably, the soluble oligomers were also not formed when mPrP was bound by the non-toxic POM2. Upon addition of scPOM-bi to mPrP, instead, the presence of soluble oligomers are detected at the first observed time point (~5 minutes after complex formation) but these disappear with time (Fig 4A).

It was previously shown that POM1-induced toxicity is inhibited by the prior incubation of PrP with POM2 [15]. Similarly, soluble oligomers were not formed when POM1 was added to a pre-formed POM2:mPrP complex (Fig 4A). By contrast, addition of POM2 to pre-formed POM1:mPrP complexes was not capable of preventing toxicity or eliminate the presence of soluble oligomers (Fig 4A).

In contrast to POM2, scPOM-bi was able to eliminate soluble oligomers even when added 5 minutes after the formation of a POM1:mPrP complex, when DLS showed that soluble oligomers were already present (Fig 4A). The difference was not due to the presence of two binding sites in scPOM-bi since the bivalent IgG versions of POM1 and POM2 behaved like their single chain counterpart.

Since DLS can only detect the presence of species in solution, we used PAGE/Western blot quantification to investigate the presence of insoluble aggregates. After formation of the mPrP:Ab complexes, part of the sample was analyzed with DLS and the remaining was subjected to 5’ centrifugation at 20’000 x g. The resulting supernatant was analyzed with PAGE to quantify the amount of mPrP and Ab present in solution (Fig. 4B). mPrP or antibodies alone neither formed aggregates over the observed time course (up to 1h, DLS analysis) nor precipitated. When the toxic POM1 antibody bound mPrP or ΔmPrP_90-230_, ~40% of the amount of mPrP and antibody was detected in the soluble fraction after centrifugation (Fig 4B); however oligomers were not formed with the truncated form of PrP, which does not cause neurotoxicity when bound to POM1. Similar levels were detected when POM2 was added after formation of the POM1:PrP complex, which is known to cause cell toxicity.

By contrast, little to no mPrP or antibody was present in solution in the non-toxic combinations such as the scPOM-bi and POM2 complexes or when POM2 was added before POM1. In contrast to POM2, however, the bispecific was able to eliminate the POM1 induced soluble oligomers when added after formation of the POM1:mPrP complex. Again, the difference was not due to the presence of two binding sites in scPOM-bi since the full IgG versions of POM1 and POM2 behaved like their single chain counterpart.

In order to further characterize oligomers and insoluble aggregates through an independent method we measured the size of mPrP:Ab complexes by confocal microscopy. Briefly, mPrP and antibody were mixed in a test tube and, after 20 minutes, analyzed by DLS to characterize the aggregation state. At that point the material was deposited on microscopy slides without centrifugation or other clarification steps. The lowest resolvable structure in laser scanning confocal microscopy images in our experimental settings is diffraction limited at ~120nm. Objects of smaller size are detected as points but the size and diameter cannot be properly resolved or measured. The toxic POM1:mPrP complexes formed smaller particles with average area of ~2.5 μm^2^. Statistically larger particles with an average of ~6 μm^2^ were detected for the non-toxic POM2 and scPOM-bi combinations (Fig 4C).

Furthermore, treatment of the mPrP:Ab complexes formed *in vitro* with 2μg/mL of Proteinase K showed that species resistant to low PK concentrations were more abundant when mPrP was bound by the toxic POM1 rather than POM2, scPOM-bi or when POM1 bound a PrP construct lacking the FT (Fig 4D).

Overall, the above indicates that protective antibodies prevented the formation of soluble, partially PK resistant oligomers whose formation requires the FT and is correlated with POM1 induced neurotoxicity. The protective scPOM-bi bispecific antibody, but not POM2, was capable of eliminating pre-formed POM1-induced soluble oligomers.

In order to test if the POM1-dependent soluble oligomers are indeed toxic, we first formed them *in vitro* by mixing mPrP with the various antibodies and then added the material to CAD5 cells; after 48 hours the cellular toxicity was evaluated with standard propidium iodide staining and FACS analysis (Fig 5). The scPOM1:mPrP soluble oligomers caused toxicity in PrP^C^-expressing CAD5 cells but not in PrP^C^ knock-out (PrP^−/−^) CAD5. No toxicity was detected, instead, when scPOM1 was in complex with the truncated ΔmPrP_90-230_ lacking the FT. Addition of the scPOM2:mPrP complex resulted in no toxicity, as well. Intriguingly, addition of the scPOM-bi:mPrP material to cells 10 minutes after complex formation resulted in toxicity, albeit lower than with scPOM1.

**Fig. 5:**
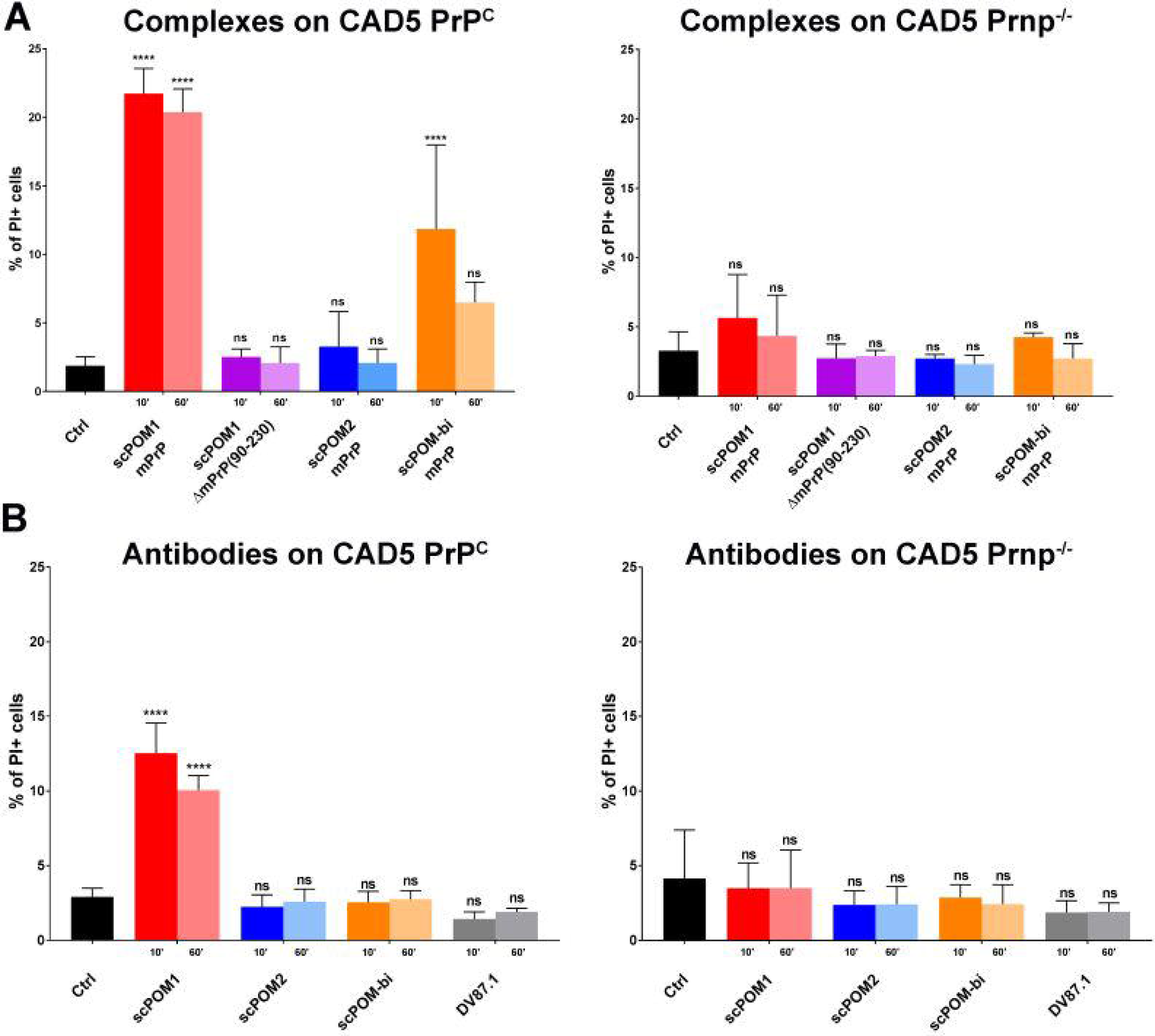
scPOM-bi does not induce toxicity on CAD5 expressing PrPC in comparison to scPOM1. The percentage of PI positive cells for different mPrP:antibodies complexes (A) or antibodies alone (B) on CAD5 PrP^C^ (left) and on CAD5 Prnp^−/−^ (right) are shown; each sample was added to cells after 10’ or 60’ of incubation at RT. The scPOM1:mPrP soluble oligomers caused toxicity in PrP^C^-expressing CAD5 cells but not in PrP^C^ knock-out (PrP^−/−^) CAD5. No toxicity was detected when scPOM1 was in complex with the truncated ΔmPrP_90-230_ lacking the FT. Addition of the scPOM2:mPrP complex resulted in no toxicity, as well. Addition of the scPOM-bi:mPrP material to cells 10 minutes after complex formation resulted in toxicity, albeit lower than with scPOM1. However, toxicity was not significant if the material was added 60 minutes after complex formation. (n=4 for all samples tested).

However, toxicity was not significant if the material was added 60 minutes after complex formation. DLS shows that scPOM-bi forms soluble oligomers, much like POM1, in the initial minutes after complex formation (possibly because some of the bispecific engages mPrP only with the POM1 moiety, thus acting just like POM1) but these disappear with time. There is, in other words, correlation between the presence of soluble oligomers and cellular toxicity. By contrast, addition of the isolated scPOM-bi to CAD5 cells resulted in no toxicity, differently from POM1. A possible interpretation is that formation of toxic scPOM-bi soluble oligomers requires local high concentration of mPrP which is available *in vitro* but not in cells.

## Discussion

The discovery that anti-PrP antibodies can block prion pathogenesis *in vivo* [30] has suggested that such antibodies might be exploited as therapeutics against human prion diseases. However, caution must be exercised as some antibodies directed against the GD of PrP can trigger PrP-mediated cellular neurotoxicity in the absence of prions [12,31]. This finding has far-reaching implications, including the possibility that autoimmunity to PrP may lead to neurodegenerative diseases. At the molecular level, it suggests that binding to specific sites in the GD can trigger changes in PrP leading to toxicity.

POM1, an antibody binding to the α1-α3 region of the GD, is acutely neurotoxic [11,17,31,32] and is thought to phenocopy the binding of PrP^Sc^ or other factors to a toxicity-triggering site of PrP^C^. According to this hypothesis, occupation of the POM1 docking site on PrP may be beneficial against prion diseases. We thus produced a bispecific antibody (termed scPOM-bi) by fusing the single chain versions of POM1, with the intent of blocking the toxicity triggering site in the GD, and POM2, which binds the FT and prevents POM1-induced toxicity.

Indeed, scPOM-bi not only lacked neurotoxicity, despite containing the toxic POM1 moiety, and in contrast to simultaneous addition of POM1 and POM2 to COCS, but also protected organotypic brain cultures from prion-induced neurodegeneration. Neuroprotection was evident even when scPOM-bi was administered 21 days post infection, when signs of prion pathology were already evident, suggesting that molecular tweezers may form the basis for a rational therapy of prion and possibly other neurodegenerative diseases. scPOM-bi afforded stronger neuroprotection than POM2, suggesting that the concomitant blockade of GD and FT is particularly effective against prion toxicity.

POM1, but not POM2 or scPOM-bi, caused recombinant mPrP to form soluble oligomers resistant to low concentration of proteinase K. Soluble oligomers were not formed when POM1 bound to an N-terminally truncated construct lacking the FT, ΔmPrP_90-230_. There is a correlation between oligomer formation and toxicity: neurotoxic antibodies or combinations (such as POM1) formed soluble oligomers, whereas those that were unable to generate them (such as POM2 or the bispecific tweezer described here) were innocuous or even protective in *ex vivo* cultured brain slices. Toxicity was detected when the POM1 induced soluble oligomers were formed in vitro from isolated, recombinant mPrP and antibody and subsequently added to PrP-expressing CAD5 cells. There was no toxicity, instead, in knock-out CAD5 lacking PrP or upon addition of the complexes formed by the protective antibodies. The bispecific is peculiar: soluble oligomers were detected by DLS 10 minutes after formation of the scPOM-bi:mPrP complex, although less abundant than those formed by POM1. Toxicity, lower than with POM1, was detected upon addition of these species to CAD5 cells. By contrast, DLS showed that scPOM-bi soluble oligomers disappear over time just like toxicity disappeared if we waited 60 minutes before adding the scPOM-bi:mPrP complex to CAD5 cells. Curiously, no toxicity was detected if scPOM-bi alone was added to CAD5 cells, whereas POM1 is toxic in this condition. A plausible explanation is that transient, soluble oligomers are only formed if scPOM-bi and mPrP are both present at high concentration (10μM in our assay), whereas the local PrP concentration in CAD5 cells is presumably lower. It is also worth noting that treatment with isolated POM1 causes apparently lower toxicity than treatment with the same amount of POM1:PrP soluble oligomers, further suggesting that the oligomers are relevant for toxicity. The above observations indicate that the induction of oligomeric PrP forms may play a role in POM1 toxicity. Since the antibodies that prevent oligomer formation were protective not only against POM1 but also against prion infection, we suggest that these soluble oligomers might also be involved in prion mediated toxicity. This interpretation is in agreement with the conjecture that soluble aggregates are the primary toxic species driving diverse proteinopathies such as Alzheimer’s and Parkinson’s disease. scPOM-bi and other prion protective antibodies may steer PrP from a toxic oligomerization to a non-toxic aggregation pathway. The fact that formation of PK resistant material is not inhibited by scPOM-bi further corroborates the idea that toxicity, or protection, is mediated by a different downstream event. There is evidence for similar mechanisms in alpha-synuclein and Aβ, where non-toxic aggregates of size and conformation different from those of toxic soluble oligomers were found [33–35]. Distinct tauopathies are linked to different molecular conformers of aggregated Tau, as well [36]. Furthermore, PrP^Sc^ conformers of different size, compatible with what we observed by DLS, and shape were found in strains with different properties and infectivity[37,38].

The scPOM-bi bispecific antibody, but not POM2, achieved the elimination of existing oligomers from a solution of pre-formed POM1:mPrP complexes. scPOM-bi was also more effective than POM2 at blocking prion mediated toxicity. One possibility is that, by bridging across two PrP molecules, scPOM-bi might favor the elongation of preexisting soluble oligomers, leading to larger, non-toxic species (Fig 6). However POM2-IgG can also bridge across two PrP molecules and yet it is less protective than scPOM-bi and cannot eliminate the POM1-induced soluble aggregates despite comparably high affinity. Engagement of the globular domain appears to be important, either by inhibiting the binding of other molecules to a toxicity triggering site or by locking PrP GD in a non-toxic conformation.

**Fig. 6:**
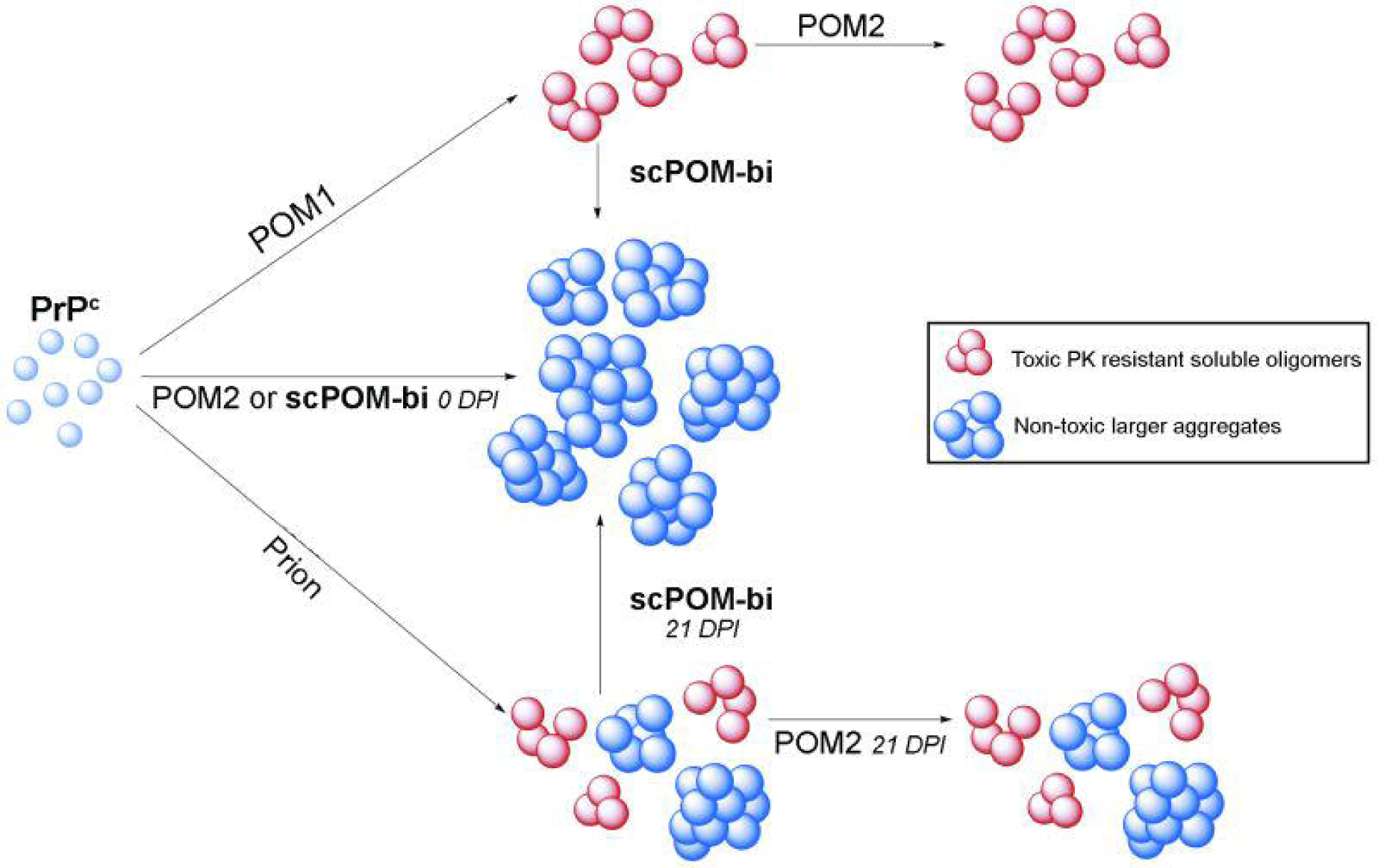
The bispeficic antibody scPOM-bi prevents the formation of soluble PrP oligomers and protects from prion neurotoxicity even when administered late after infection. Addition of POM1 antibody (top), but not POM2 or scPOM-bi, to PrP^C^ generates soluble, pK resistant PrP oligomers (red) whose presence correlates with toxicity (top); subsequent addition of the neuroprotective scPOM-bi, but not POM2, eliminates them in favor of larger, non-toxic aggregates (blue). Small soluble oligomers might also be responsible for prion induced toxicity (bottom) similarly to other amyloidosis. scPOM-bi might be able to eliminate them just as it does with the POM1-induced oligomers whereas POM2 might not, which would explain why only the bispecific is neuroprotective even at late administration (21 days post infection, dpi).

Indeed, simultaneous targeting of GD and FT by the bispecific antibody described here resulted in more potent protection, even when given at late timepoints, than simple targeting of the FT. We conclude that the strategy delineated here may be exploited for the development of effective immunotherapeutics against prion and possibly other diseases.

## Materials and Methods

### Cerebellar organotypic slice cultures (COCS)

Preparation of COCS was undertaken as described elsewhere [39]. Briefly, 350 µm thick COCS were prepared from 9-12 day old Tga20 or ZH3 pups [40]. Inoculation of COCS was performed with 100 µg brain homogenate per 10 slices from terminally sick prion-infected (RML6) or NBH from CD1 mice, diluted in 2 mL physiological Grey’s balanced salt solution [41]. The infectious brain homogenate was added to the free-floating COCS for 1 h at 4°C then washed, and 5–10 slices were placed on a 6-well PTFE membrane insert. Antibodies were first added 1, 10 or 21 days post-infection then re-added with every medium change (three times a week).

### CAD5 PrP^C^ and Prnp^−/−^

In order to assess PrP^C^-dependent toxicity of the soluble oligomers *in vitro* we generated *Prnp* knock-out versions (*Prnp^−/−^*) of the murine neuroblastoma cell line CAD5 by CRISPR/Cas9. CAD5 is a subclone of the central nervous system catecholaminergic cell line CAD showing particular susceptibility to prion infection [38,42]. To avoid expression of abberant or truncated versions of PrP^C^ or deletion of the splice acceptor site that would lead to pathological overexpression of Doppel (Dpl) mRNA [43], we used single-stranded guide RNA (sgRNA) cloned into the MLM3636 plasmid aiming at a protospacer adjacent motif (PAM) site in the signal peptide of *Prnp* (S7A Fig.). Cells were co-transfected with MLM3636 and the hCas9 plasmid followed by transient antibiotic selection.

Subsequently, 7 clones were manually picked, expanded and subjected to further characterization. Cells were lysed and PrP^C^ levels were measured by POM1/POM2 sandwich ELISA. Herein, 7 CAD5 *Prnp*^−/−^ candidate clones #A6, #C2, #C6, #C12, #H7, #H9 and #H12 all showed near-zero PrP^C^ levels comparable to the established *Prnp*^−/−^ cell line HPL (p>0.05, one-way ANOVA with Dunnett’s post-hoc test, S7B Fig.) [44]. 5 clones were further assessed on western blot, where no detectable PrP^C^ levels could be observed (S7C Fig.), suggesting a successful knock-out of PrP^C^ in all 5 *Prnp*^−/−^ candidate clones. DNA was extracted from expanded cells of clones #C2 and #C12 and the mutagenized region was sequenced by PCR amplification of the open reading frame (ORF) of *Prnp*. Amplified products were cloned into the pCR-Blunt II-TOPO plasmid (Invitrogen) and 10 colonies per clone were sequenced by Sanger sequencing. Herein, #C2 showed four different mutations, i.e., three deletions and one insertion, while #C12 showed two different deletions proximal to the PAM (S7D Fig.). These results indicate multiploidy to be more likely in #C2 than in #C12, hence all further experiments are performed with the CAD5 *Prnp*^−/−^ clone #C12.

For CRISPR/Cas9-aided generation of CAD5 knock-out cells, mouse *Prnp* sgRNA was designed using the web-based tools http://crispr.mit.edu/ and http://zifit.partners.org/ZiFiT/CSquare9GetOligos.aspx (last access on May 15^th^ 2017). The sgRNA expression plasmid MLM3636 was a gift from Keith Joung (Addgene plasmid # 43860, www.addgene.org). For annealing of single-stranded DNA oligomers of sgRNA for subsequent cloning into the MLM3636 plasmid the following ligation reaction was prepared: 10 µl Oligo4 F [100 µM] (5’ - ACA CCG CAG TCA TCA TGG CGA ACC TG - 3’), 10 µl Oligo4 R [100 µM] (5’ - AAA ACA GGT TCG CCA TGA TGA CTG CG - 3’), 10 µL of NEB Buffer 2.1 (New England Biolabs), 70 µl ddH_2_O. Reaction mix was heated for 4 min at 95°C on a heating block ThermoStat (Eppendorf), then the heating block was turned off and the reaction was allowed to proceed for 30 min on the block and was then put at 4°C. Golden Gate assembly [45] was used in order to clone the double-stranded DNA Oligomers into the MLM3636 plasmid, using the following reaction:

| *amount* | *name* |
| --- | --- |
| 150 ng | MLM3636 plasmid |
| 1 µl | double-stranded oligomer ligation mix |
| 2 µl | NEB T4 ligase buffer (New England Biolabs) |
| 13.25 µl | ddH<sub>2</sub>O |
| 1 µl | Esp3I (New England Biolabs) |
| 1 µl | T4 ligase (New England Biolabs) |

This reaction was put on a thermocycler using the following conditions:

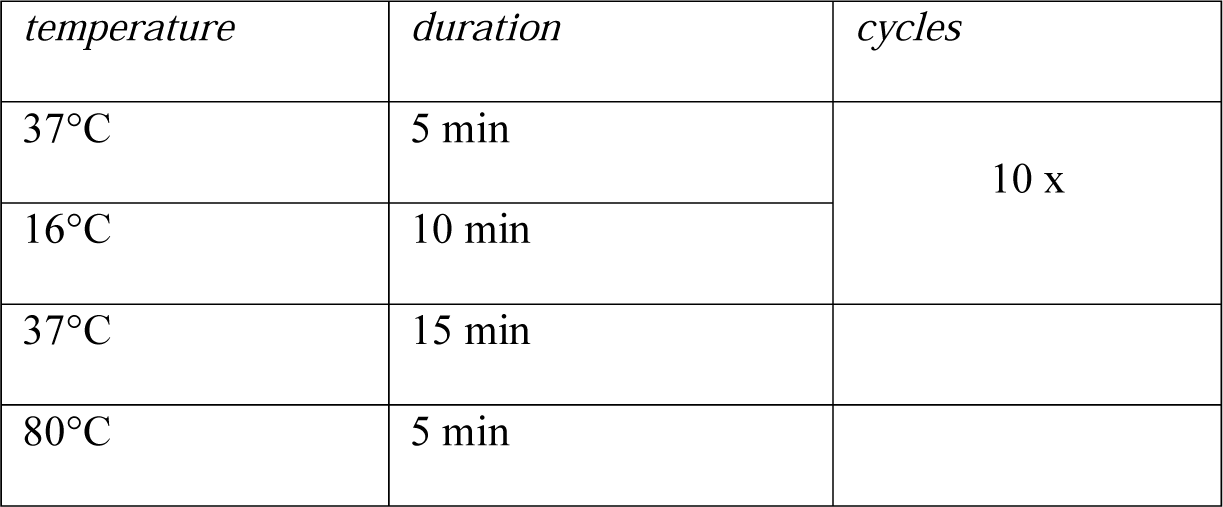

the double-stranded DNA Oligomers into the MLM3636 plasmid, using the following reaction: This reaction was put on a thermocycler using the following conditions:
| *temperature* | *duration* | *cycles* |
| --- | --- | --- |
| 37°C | 5 min | 10 x |
| 16°C | 10 min |  |
| 37°C | 15 min |  |
| 80°C | 5 min |  |

The ligated plasmid MLM3636(sgRNA_m*Prnp*_) was subsequently transformed into DH5α chemically competent *E. coli* cells (Invitrogen) and plasmid purification was undertaken using Plasmid Maxi Kit (Qiagen). CAD5 cells were co-transfected using the MLM3636(sgRNA_m*Prnp*_) plasmid and the hCas9 plasmid (hCas9 was a gift from George Church, Addgene plasmid # 41815, [46]) dissolved in Lipofectamine 2000 (Invitrogen). After selection of transfected cells with Geneticin (Invitrogen), single colonies were picked and expanded. For sequencing, DNA was extracted from cells using DNeasy Blood & Tissue Kit (Qiagen). PCR amplification with Q5 high-fidelity DNA polymerase was undertaken using the primers Prn-ko F1 (5’ - TGC AGG TGA CTT TCT GCA TTC TGG - 3’) and P10 rev (5’ - GCT GGG CTT GTT CCA CTG ATT ATG GGT AC - 3’). After PCR clean-up using NucleoSpin Gel and PCR Clean-up kit (Macherey-Nagel), blunt-end PCR fragments were cloned into Zero Blunt TOPO PCR Cloning Kit (Thermo Fisher Scientific) and Sanger Sequencing (Microsynth) was performed to identify mutated *Prnp* sequences. Western Blot and ELISA was undertaken to confirm *Prnp*^−/−^ as described below. Unless mentioned otherwise, clone #C12 was used for all experiments.

### Enzyme-linked immunosorbent assay (ELISA)

For measuring PrP^C^ levels from cell lysates, ELISA was performed as described previously [32]. Herein, 96-well MaxiSorp polystyrene plates (Nunc) were coated with 400 ng/ml POM1 (or POM19) in PBS at 4°C overnight. Plates were washed three times in PBS + 0.1% Tween-20 (0.1% PBS-T) and blocked with 80 µl per well of 5% skim milk in 0.1% PBS-T for 1.5 h at room temperature. Blocking buffer was discarded and samples and controls were added dissolved in 1% skim milk in 0.1% PBS-T for 1 h at 37°C. Recombinant, full-length murine PrP^C^ (rmPrP_23-230_) was used as positive control, 0.1% PBS-T was used as negative control. Biotinylated POM2 was used to detect PrP^C^ (200 ng/ml in 1% skim milk in 0.1% PBS-T).

### CAD5 toxicity assay

Quantification of toxicity on CAD5 either expressing or lacking PrP^C^ induced by different antibodies alone or in complex with mPrP/ΔmPrP(90-230) was measured as percentage of PI positive cells using Flow Cytometry. CAD5 PrP^C^ or Prnp^−/−^ were cultured with 20mL Corning™ Basal Cell Culture Liquid Media - DMEM and Ham’s F-12, 50/50 Mix supplemented with 10% FBS, Gibco™ MEM Non-Essential Amino Acids Solution 1X, Gibco™ GlutaMAX™ Supplement 1X and 0,5mg/mL of Geneticin in T75 Flasks ThermoFisher™ at 37*C 5% CO_2_. 16 hours before treatment, cells were splitted into 96wells plates at 25000cells/well in 100μL.

Complexes of PrP:Antibodies (1:1 PrP:Ab ratio for monovalent binding, 2:1 PrP:Ab ratio for bivalent binding) or antibodies alone were formed at 5μM final concentration, in 20mM HEPES pH 7,2 150mM NaCl. After 10’ or 60’ upon complex formation, 100μL of each sample, including buffer alone or unrelated antibodies as controls, were added to CAD5 cells, in duplicates. After 48 hours, cells were washed two times with 100μL MACS buffer (PBS + 1% FBS + 2mM EDTA) and resuspended in 100μL MACS buffer. 30” before FACS measurements PI (1μg/mL) was added to cells. Measurements were performed using BD™ LSRFORTESSA Statistics: percentage of PI positive cells were plotted in columns as mean with SD. 2way ANOVA test with Tukey test was performed comparing each samples (* p<0.05, ** p<0.005, *** p<0.0005, **** p<0.00001)

### Immunohistochemical stainings and NeuN morphometry

45 days post infection, prior to fixation, COCS were rinsed twice in PBS, fixed in 4% paraformaldehyde for 2 days at 4°C, washed twice in PBS and incubated in blocking buffer (0.05% Triton X-100 vol/vol, 0.3% goat serum vol/vol in PBS) for 1 hour at room temperature. NeuN stainings were performed for 3 days at 4°C with an Alexa-488 conjugated mouse anti-NeuN antibody (Life Technologies) at 1.6 µg/mL in blocking buffer and incubated. After washing for 15 min in PBS, cell nuclei were made visible by a 30 min incubation with DAPI (1 µg/mL) in PBS at room temperature. Slices were then washed again twice in PBS for 15 minutes and mounted with fluorescent mounting medium (DAKO) on a glass slide. NeuN morphometry of COCS was undertaken on images taken on a fluorescent microscope (BX-61, Olympus) at identical exposure times through custom written scripts for the image analysis software cell^P (Olympus) as previously described [11].

### Proteinase K digestion (COCS)

COCS were washed twice in PBS and scraped off the slice culture membrane using 10 µL PBS per slice. Slice cultures were homogenized using a TissueLyser LT (Qiagen) small bead mill at 50 Hz for 2 minutes. For determination of PrP^Sc^, RML6 (1 µl of 10% w/v brain homogenate or 2 µg protein per lane) and slice culture homogenates (20 µg protein per lane) of *Tga20* COCS were digested with 25 μg mL-^1^ proteinase K (Roche) at a final volume of 20 μL in PBS for 30 minutes at 37°C. For inactivation of proteinase K, 7 µL of 4x NuPAGE LDS sample buffer (Thermo Fisher Scientific) was added and samples were boiled at 95°C for 5 minutes. Equal sample volumes were loaded on Nu-PAGE Bis/Tris precast gels (Life Technologies) and Western blotting was performed using the monoclonal anti-PrP antibody POM1 as described elsewhere[32].

### Computational Modelling and Molecular Dynamics of scPOM-bi

The structure of scPOM-bi used in the computational simulations was modeled on the experimental structures of the POM1:PrP_124-225_ complex (PDB code 4H88) and POM2:octarepeat-peptide (PDB code: 4J8R) complexes. After initial superposition of the POM1 and POM2 moiety on the structure of one (for intramolecular binding models) or two (for intermolecular binding models) PrP molecules, the linker joining the two scFv was manually built. The system was then subjected to 100ns of fully atomistic molecular dynamics simulations (MD) to obtain an energetically favorable and stable conformation.

In all cases, the system was initially set up and equilibrated through standard MD protocols: proteins were centered in a triclinic box, 0.2nm from the edge, filled with SPCE water model and 0.15M Na+Cl- ions using the AMBER99SB-ILDN protein force field. Energy minimization was performed in order to let the ions achieve a stable conformation. Temperature and pressure equilibration steps, respectively at 298°K and 1 Bar, of 100ps each were then performed before performing 300ns molecular dynamics simulations with the above mentioned force field. MD trajectory files were analyzed after removal of Periodic Boundary Conditions. The stability of each simulated complex was verified by root mean square deviation, radius of gyration and visual analysis.

### Protein production

Recombinant mouse PrP full length (23-230), PrP lacking FT (90-230), globular domain only (120-230) or Flexible tail only (23-120) were expressed and purified from *E.coli* as previously described[47,48]. scPOM1, scPOM2 and scPOM-bi sequences were codon optimized, cloned in frame into a pET21a plasmid (Novagen) and expressed in *E.coli* Rosetta (scPOM-bi) or Rosetta pLysS (for scPOM1 and scPOM2) cells. Bacterial cells were grown in 2XYT media plus Ampicillin (100µg/mL) and chloramphenicol (34µg/mL) at 25°C (scPOM1 and scPOM2) or 37°C (scPOM-bi) until OD600 reached 0.6, then induced with 0.5mM IPTG and harvested 16 (scPOM1 and scPOM2) or 3 (scPOM-bi) hours post-induction. Bacterial pellets were sonicated with 50mM MES pH6.5 100mM NaCl and 0.5mM DTT (50mL per liter of colture) and centrifuged at 16500rpm (rotor ss34) for 30’ at 4°C. The pellet was then washed with sonication buffer plus 0.5% of Triton-X100 and then with sonication buffer to remove excess of Triton-X100. The pellet was solubilized in 50mM MES pH 6.5 1M NaCl 6M Guanidine-HCl (Buffer A). Following addition of 0.2% PEI and centrifugation to remove DNA contamination, 60% ammonium sulfate was added to the supernatant to remove traces of PEI. After centrifugation the pellet was resuspended in Buffer A, filtered and loaded on pre-equilibrated HisTrap Column (GE healthcare) with Buffer A. The column was washed with at least 5 volumes of Buffer A and eluted with Buffer A plus 500mM Imidazole. Antibody containing fractions were diluted 20 times into refolding buffer (20 mM phosphate buffer pH 10.5, 100 mM NaCl, 200 mM arginine, 1mM Glutathione Reduced and 0.1mM Glutathione Oxidized). scPOM1, scPOM2 and scPOM-bi were finally purified on a Superdex-75 size exclusion column (GE) in PBS buffer. The elution and dynamic light scattering profiles of all proteins were consistent with monomeric species. Full IgG POM1 and POM2 monoclonal antibodies were produced and purified as described previously[32].

### SPR binding assays

The binding properties of the complexes between scPOM1, scPOM2, either in single chain or IgG version, and scPOM-bi on different mPrP constructs were analyzed at 25°C on a ProteOn XPR-36 instrument (Bio-Rad) using 20mM HEPES pH 7.2 150mM NaCl 3mM EDTA and 0.005% Tween-20 as running buffer. 50nM solutions of PrP constructs were immobilized on the surface of a GLC sensor chip through standard amine coupling. Serial dilution of antibodies in the nanomolar range were injected at a flow rate of 100 µL/min (contact time 6 minutes); dissociation was followed for 5 minutes. Analyte responses were corrected for unspecific binding and buffer responses by subtracting the signal of both a channel were no PrP was immobilized and a channel were no antibody was added. Curve fitting and data analysis were performed with Bio-Rad ProteOn Manager software (version 3.1.0.6). In the PrP dilution experiments used to evaluate avidity effects, serial dilution (1:100, 1:200, 1:500 and 1:1000) of NHS/EDAC compounds were used for GLC chip surface activation in order to limit the amount of immobilized protein.

### Dynamic Light Scattering (DLS) assays

The size of PrP either alone or in complex with different antibodies was estimated by Dynamic Light Scattering (DLS) at 25°C using a “DynaPro - NanoStar” instrument (WYATT) in 10µL at a concentration of 5µM. The PrP:antibody ratio was 1:1 for monovalent binding (e.g. scF_v_) and 2:1 for bivalent binding species (e.g. full IgG). Readings were taken 2, 5, 10, 20 and 30 minutes after complex formation. When evaluating addition of a second antibody (e.g. forming a PrP/scPOM1 complex and then adding scPOM-bi), PrP and the first antibody were pre-mixed for 5 minutes and then the second antibody was added. Each time point measurement was performed by 10 repetitions of 5 seconds signal integration. A Rayleigh sphere model was used for size estimation. At least 3 repetitions of the same experiment were performed on different, freshly prepared samples. Before complex formation, all samples were dialyzed together in 20mM HEPES pH 7.2 150mM NaCl, centrifuged at 21000g and filtered with 0.22µm filters before measurement. Statistics: all experiments are shown as mean with standard error of the mean (SEM).

### Precipitation assays

Quantification of soluble PrP either alone or in complex with different antibodies was performed using SDS-PAGE and either comassie staining (for PrP alone and in complex with scFv) or western blot (for PrP/IgG complexes). The precipitation assays were run in parallel to DLS assays in the conditions indicated above. Samples were centrifuged for 2 minutes at 21000g at 4°C. 25µL of supernatant was collected, mixed with equal volume of sample buffer and loaded on polyacrylamide gel (4% Stacking – 12% running). For comassie staining, SDS-PAGEs were left 10 minutes in 2.5g/L Comassie Brilliant Blue G-250 (Sigma) 40% Methanol and 10% Glacial Acetic Acid and then destained using 70% ddH20, 20% Methanol and 10% Glacial Acetic Acid for 1hour at least.

Gels were then acquired using ImageQuant LAS 4000 (GE Healthcare) according to standard procedures. For western blot, proteins from SDS-PAGE were transferred onto PVDF membranes, blocked in TBS-Tween20 10% Milk for 10 minutes at RT and probed with an antibody against PrP (POM19 mouse IgG 1µg/mL in TBS-Tween20 for 16 hours at 4°C) that does not compete with either scPOM1, scPOM2 and scPOM-bi. The primary antibody was detected using a goat anti-Mouse-HRP conjugated antibody (1:10000 in TBS-Tween20 for 1hour at RT, from ThermoFisher) and developed using Pierce™ ECL Western Blotting Substrate (ThermoFisher).

Chemiluminescence from PrP specific bands were acquired using a ImageQuant LAS 4000 (GE Healthcare) using High Resolution for sensitivity, 1/60 or 1/100 sec exposure time and Precision as exposure type. Quantification of PrP was then performed using Multi Gauge Software (from FujiFilm) with standard protocol, normalizing all samples to PrP Alone control bands. At least 3 independent replicates of the experiments were performed. Statistics: all experiments are shown as mean with SEM.

### Confocal analyses of PrP/Ab Oligomers

Antibodies were labelled with Alexa Fluor™ 647 NHS Ester (Thermo Fisher Scientific) in PBS carbonate pH 8.3 with a 1:2 Ab:Dye ratio; unbound dyes were removed using Size Exclusion Chromatography. Complexes between mPrP and Abs were generated at 5μM in 10μL with 1:1 ratio for monovalent binding (e.g. scF_v_) and 2:1 for bivalent binding species (e.g. POM-bi) as for DLS assays. After 10’ of incubation at 25C, 2μL of each complex was added to glass microscopy slides (Thermo Fisher Scientific) and covered with coverslip. The same samples were also subjected to DLS analyses, in parallel. Images were acquired using a Leica TCS SP5 confocal microscope using sequential acquisition settings to visualize aggregates containing labelled antibodies. For each mPrP:Ab complex, 4 fields of view of 246 μm x 246 μm were acquired with a 63X/1.4 NA oil immersion objective. Images were analysed using IMARIS software (Bitplane). To estimate particles size surfaces were generated in software based on the fluorescent signal from Alexa647 dye (segmentation parameters: surface grain size 0.01 μm, intensity threshold set at 50). Statistics: all particles were shown in dot plot graph as mean with SD. Mann-Whitney test was performed (* p<0.05, ** p<0.01, *** p<0.001, **** p<0.0001)

### Ethics Statement

All animal experiments were conducted in strict accordance with the Rules and Regulations for the Protection of Animal Rights (Tierschutzgesetz and Tierschutzverordnung) of the Swiss Bundesamt für Lebensmittelsicherheit und Veterinärwesen BLV. Body weights were measured weekly. All animal protocols and experiments performed were specifically approved for this study by the responsible institutional animal care committee, namely the Animal Welfare Committee of the Canton of Zürich (permit numbersVersuchstierhaltung 123, ZH139/16 and 90/2013). All efforts were made to minimize animal discomfort and suffering.

## Acknowledgments

We thank Robyn Grace Holden for graphic assistance and Diego Morone for microscope technical support.

## Supporting information

**S1 Text:** SPR analysis of scPOM-bi binding to PrP

**S1 Movie:** fully atomistic molecular dynamics simulation of 30 ns of the model of scPOM-bi in complex with one molecule mPrP.

**S2 Movie:** fully atomistic molecular dynamics simulation of 30 ns of the model of scPOM-bi in complex with two molecules mPrP.

**S1 Fig:** thermal denaturation of scPOM-bi measured by CD spectroscopy, indicating a melting temperature of 75°C

**S2 Fig:** Fluorescent micrographs of all biological replicates (A, scPOM-bi; B, POM2 IgG). Scale bar = 500 µm.

**S3 Fig:** values of association (k_a_, left) and dissociation constants (k_d_, right) for scPOM1 (yellow), POM1-IgG (red) and scPOM-bi (blue) at different concentrations of PrP, measured by SPR. The dilution of PrP on the sensor chip is reported. The dissociation constant, but not the association, is affected by PrP dilution, indicating that intermolecular avidity effects are present in POM1 IgG and scPOM-bi. See S1 Text for further details.

**S4 Fig:** Representative confocal microscopy images of PrP:Ab oligomers and aggregates (scale bar = 10μm). See main text (Fig 4) for quantification and methods for experimental details. Briefly, complexes between recombinant mPrP and antibodies were formed in vitro and the material deposited on microscopy slides without centrifugation or other purification steps. Species of different size are apparent when mPrP is in complex with toxic (POM1) or non toxic antibodies.

**S5 Fig:** Precipitation assays confirmed that the toxic scPOM1:mPrP complex generates soluble oligomers containing both PrP and antibody. No soluble material was present in the complexes between mPrP and scPOM2, scPOM-bi or when scPOM-bi was added after scPOM1. See main text (Fig 4) for quantification and methods for experimental details. Briefly, after formation of the mPrP:Ab complexes *in vitro* the samples were centrifuged at 20’000 x g. The amount of mPrP and Ab present in the resulting supernatant was estimated with PAGE/Western Blot. The quantity of soluble material is reported as percentage of soluble mPrP or Ab alone, which do not precipitate or form aggregates over the observed time frame. The variability is due to the experimental set up and to the fact that we are analyzing transient, non-homogeneous species that are likely to change over time. Such variability does not affect the statistical significance of the measures.

**S6 Fig:** low concentration PK resistance assay. mPrP:Ab complexes were formed *in vitro* and 2μg/mL of Proteinase K were added. The presence of PK resistance species was assessed by western blot. An increased amount of species resistant to PK was detected in the scPOM1:mPrP complexes, but only if the flexible tail was present, which correlates to toxicity and protection assays. See main text (Fig 4) for quantification and statistics.

**S7 Fig:** Generation of a stable CAD5 Prnp^−/−^ cell line. (A) Design of sgRNA for CRISPR/Cas9 mediated generation of CAD5 *Prnp*^−/−^ cells. A PAM in the coding sequence of the signal peptide was chosen. (B) ELISA of 7 candidate CAD5 *Prnp*^−/−^ clones showed similar PrP^C^ levels compared to the established *Prnp*^−/−^ cell line HPL (p>0.05, one-way ANOVA with Dunnett’s post-hoc test, all clones versus HPL), 5 of which were further assessed by PrP^C^ western blot, confirming lack of PrP^C^ expression (C). (D) Sanger sequencing of PCR amplified *Prnp* ORF showed n=4 different mutations in #C2 and n=2 different mutations, labelling according to (A). The splice acceptor site is unaffected in both of the constructs.

## Reference

1. Aguzzi A, Sigurdson C, Heikenwaelder M (2008) Molecular mechanisms of prion pathogenesis. Annu Rev Pathol 3: 11–40.

2. Aguzzi A, Calella AM (2009) Prions: protein aggregation and infectious diseases. Physiol Rev 89: 1105–1152.

3. Aguzzi A, Lakkaraju AK (2016) Cell Biology of Prions and Prionoids: A Status Report. Trends Cell Biol 26: 40–51.

4. Waddell L, Greig J, Mascarenhas M, Otten A, Corrin T, et al. (2018) Current evidence on the transmissibility of chronic wasting disease prions to humans-A systematic review. Transbound Emerg Dis 65: 37–49.

5. Dyrbye H, Broholm H, Dziegiel MH, Laursen H (2008) The M129V polymorphism of codon 129 in the prion gene (PRNP) in the Danish population. Eur J Epidemiol 23: 23–27.

6. Mok T, Jaunmuktane Z, Joiner S, Campbell T, Morgan C, et al. (2017) Variant Creutzfeldt-Jakob Disease in a Patient with Heterozygosity at PRNP Codon 129. N Engl J Med 376: 292–294.

7. Aguzzi A, Polymenidou M (2004) Mammalian prion biology: one century of evolving concepts. Cell 116: 313–327.

8. Reardon S (2015) Antibody drugs for Alzheimer’s show glimmers of promise. Nature 523: 509–510.

9. Sevigny J, Chiao P, Bussiere T, Weinreb PH, Williams L, et al. (2016) The antibody aducanumab reduces Abeta plaques in Alzheimer’s disease. Nature 537: 50–56.

10. Chung E, Ji Y, Sun Y, Kascsak RJ, Kascsak RB, et al. (2010) Anti-PrPC monoclonal antibody infusion as a novel treatment for cognitive deficits in an Alzheimer’s disease model mouse. BMC Neurosci 11: 130.

11. Sonati T, Reimann RR, Falsig J, Baral PK, O’Connor T, et al. (2013) The toxicity of antiprion antibodies is mediated by the flexible tail of the prion protein. Nature 501: 102–106.

12. Solforosi L, Criado JR, McGavern DB, Wirz S, Sanchez-Alavez M, et al. (2004) Cross-linking cellular prion protein triggers neuronal apoptosis in vivo. Science 303: 1514–1516.

13. Baral PK, Wieland B, Swayampakula M, Polymenidou M, Rahman MH, et al. (2012) Structural studies on the folded domain of the human prion protein bound to the Fab fragment of the antibody POM1. Acta Crystallogr D Biol Crystallogr 68: 1501–1512.

14. Aguzzi A, Weissmann C (1997) Prion research: the next frontiers. Nature 389: 795–798.

15. Herrmann US, Sonati T, Falsig J, Reimann RR, Dametto P, et al. (2015) Prion infections and anti-PrP antibodies trigger converging neurotoxic pathways. PLoS Pathog 11: e1004662.

16. Goniotaki D, Lakkaraju AKK, Shrivastava AN, Bakirci P, Sorce S, et al. (2017) Inhibition of group-I metabotropic glutamate receptors protects against prion toxicity. PLoS Pathog 13: e1006733.

17. Frontzek K, Pfammatter M, Sorce S, Senatore A, Schwarz P, et al. (2016) Neurotoxic Antibodies against the Prion Protein Do Not Trigger Prion Replication. PLoS One 11: e0163601.

18. Falsig J, Julius C, Margalith I, Schwarz P, Heppner FL, et al. (2008) A versatile prion replication assay in organotypic brain slices. Nat Neurosci 11: 109–117.

19. Brinkmann U, Kontermann RE (2017) The making of bispecific antibodies. MAbs 9: 182–212.

20. Mack M, Riethmuller G, Kufer P (1995) A small bispecific antibody construct expressed as a functional single-chain molecule with high tumor cell cytotoxicity. Proc Natl Acad Sci U S A 92: 7021–7025.

21. Baral PK, Wieland B, Swayampakula M, Polymenidou M, Aguzzi A, et al. (2011) Crystallization and preliminary X-ray diffraction analysis of prion protein bound to the Fab fragment of the POM1 antibody. Acta Crystallogr Sect F Struct Biol Cryst Commun 67: 1211–1213.

22. Swayampakula M, Baral PK, Aguzzi A, Kav NN, James MN (2013) The crystal structure of an octapeptide repeat of the prion protein in complex with a Fab fragment of the POM2 antibody. Protein Sci 22: 893–903.

23. Falsig J, Sonati T, Herrmann US, Saban D, Li B, et al. (2012) Prion pathogenesis is faithfully reproduced in cerebellar organotypic slice cultures. PLoS Pathog 8: e1002985.

24. Cleary JP, Walsh DM, Hofmeister JJ, Shankar GM, Kuskowski MA, et al. (2005) Natural oligomers of the amyloid-beta protein specifically disrupt cognitive function. Nat Neurosci 8: 79–84.

25. Shankar GM, Li S, Mehta TH, Garcia-Munoz A, Shepardson NE, et al. (2008) Amyloid-beta protein dimers isolated directly from Alzheimer’s brains impair synaptic plasticity and memory. Nat Med 14: 837–842.

26. Benilova I, Karran E, De Strooper B (2012) The toxic Abeta oligomer and Alzheimer’s disease: an emperor in need of clothes. Nat Neurosci 15: 349–357.

27. Ow SY, Dunstan DE (2014) A brief overview of amyloids and Alzheimer’s disease. Protein Sci 23: 1315–1331.

28. Sengupta U, Nilson AN, Kayed R (2016) The Role of Amyloid-beta Oligomers in Toxicity, Propagation, and Immunotherapy. EBioMedicine 6: 42–49.

29. Silveira JR, Raymond GJ, Hughson AG, Race RE, Sim VL, et al. (2005) The most infectious prion protein particles. Nature 437: 257–261.

30. Heppner FL, Musahl C, Arrighi I, Klein MA, Rulicke T, et al. (2001) Prevention of scrapie pathogenesis by transgenic expression of anti-prion protein antibodies. Science 294: 178–182.

31. Reimann RR, Sonati T, Hornemann S, Herrmann US, Arand M, et al. (2016) Differential Toxicity of Antibodies to the Prion Protein. PLoS Pathog 12: e1005401.

32. Polymenidou M, Moos R, Scott M, Sigurdson C, Shi YZ, et al. (2008) The POM monoclonals: a comprehensive set of antibodies to non-overlapping prion protein epitopes. PLoS One 3: e3872.

33. Lashuel HA, Overk CR, Oueslati A, Masliah E (2013) The many faces of alpha-synuclein: from structure and toxicity to therapeutic target. Nat Rev Neurosci 14: 38–48.

34. Sakono M, Zako T (2010) Amyloid oligomers: formation and toxicity of Abeta oligomers. FEBS J 277: 1348–1358.

35. Waxman EA, Giasson BI (2009) Molecular mechanisms of alpha-synuclein neurodegeneration. Biochim Biophys Acta 1792: 616–624.

36. Simic G, Babic Leko M, Wray S, Harrington C, Delalle I, et al. (2016) Tau Protein Hyperphosphorylation and Aggregation in Alzheimer’s Disease and Other Tauopathies, and Possible Neuroprotective Strategies. Biomolecules 6: 6.

37. Deleault NR, Walsh DJ, Piro JR, Wang F, Wang X, et al. (2012) Cofactor molecules maintain infectious conformation and restrict strain properties in purified prions. Proc Natl Acad Sci U S A 109: E1938–1946.

38. Mahal SP, Baker CA, Demczyk CA, Smith EW, Julius C, et al. (2007) Prion strain discrimination in cell culture: the cell panel assay. Proc Natl Acad Sci U S A 104: 20908–20913.

39. Falsig J, Aguzzi A (2008) The prion organotypic slice culture assay--POSCA. Nat Protoc 3: 555–562.

40. Fischer M, Rulicke T, Raeber A, Sailer A, Moser M, et al. (1996) Prion protein (PrP) with amino-proximal deletions restoring susceptibility of PrP knockout mice to scrapie. EMBO J 15: 1255–1264.

41. Falsig J, Julius C, Margalith I, Schwarz P, Heppner F, et al. (2008) A versatile prion replication assay in organotypic brain slices. Nat Neurosci 11: 109–117.

42. Qi Y, Wang JK, McMillian M, Chikaraishi DM (1997) Characterization of a CNS cell line, CAD, in which morphological differentiation is initiated by serum deprivation. J Neurosci 17: 1217–1225.

43. Moore RC, Lee IY, Silverman GL, Harrison PM, Strome R, et al. (1999) Ataxia in prion protein (PrP)-deficient mice is associated with upregulation of the novel PrP-like protein doppel. J Mol Biol 292: 797–817.

44. Kuwahara C, Takeuchi AM, Nishimura T, Haraguchi K, Kubosaki A, et al. (1999) Prions prevent neuronal cell-line death. Nature 400: 225–226.

45. Engler C, Kandzia R, Marillonnet S (2008) A one pot, one step, precision cloning method with high throughput capability. PLoS One 3: e3647.

46. Mali P, Yang L, Esvelt KM, Aach J, Guell M, et al. (2013) RNA-guided human genome engineering via Cas9. Science 339: 823–826.

47. Hornemann S, Christen B, von Schroetter C, Perez DR, Wuthrich K (2009) Prion protein library of recombinant constructs for structural biology. FEBS J 276: 2359–2367.

48. Zahn R, von Schroetter C, Wüthrich K (1997) Human prion proteins expressed in Escherichia coli and purified by high-affinity column refolding. FEBS Letters 417: 400–404.

